# Loss of Notch signaling disrupts ventricular wall integrity and causes obstructive hydrocephalus

**DOI:** 10.64898/2026.11.17.767352

**Authors:** Lei Gao, Lily Nguyen, Tessa Marks, Dorothy Lartey, Dhyana Magesh, Catherine E. Dyevich, Grace Siauw, Anna IL Ostrem, Caroline A. Pearson, Bennett G. Novitch, Jennifer H. Kong

## Abstract

Hydrocephalus is a common and severe neurodevelopmental disorder characterized by abnormal accumulation of cerebrospinal fluid (CSF) within the ventricular system. Although its genetic basis is complex, recent studies have linked congenital hydrocephalus to defects in neural progenitor cell (NPC) maintenance, though how this results in ventricular obstruction remains unclear. Here we show that Notch signaling not only maintains NPCs within the ventricular zone (VZ) but also preserves the structural integrity of the developing ventricular system. Conditional ablation of the Notch effector *Rbpj* elicited premature differentiation of NPCs into neurons and disrupted VZ continuity. Cells lining the VZ lost their columnar organization, apical polarity, and adhesion contacts, breaking down the neuroepithelial layer and ependymal lining. This disruption obstructed CSF flow, causing hydrocephalus in neonatal mice. Pharmacological inhibition of Notch signaling in cortical spheroids similarly disrupted neural rosette formation, recapitulating aspects of the neuroepithelial defects observed in vivo. Collectively, our findings reveal critical roles for Notch signaling in preserving the walls of the ventricular system during brain development and provide insights into how disruptions to this pathway contribute to hydrocephalus.

## INTRODUCTION

Hydrocephalus is a severe neurodevelopmental disorder characterized by the abnormal accumulation of cerebrospinal fluid (CSF) within the ventricular system, leading to increased intracranial pressure and progressive brain injury^1,2^. Globally, approximately 400,000 new cases of pediatric hydrocephalus occur each year, with more than 40% predicted to have genetic origins^3–5^. In infants, hydrocephalus is often detected through accelerated head growth and progressive macrocephaly. However, clinical manifestations are wide-ranging and can include locomotor deficits, behavioral changes, language regression, visual disturbances, headaches, and vomiting^6^. Despite its prevalence, the cellular and molecular mechanisms underlying congenital hydrocephalus remain poorly understood. Recently, whole-exome sequencing studies have identified numerous novel candidate mutations associated with congenital hydrocephalus^7–12^. While these candidate mutations disrupt diverse biological processes, many converge on pathways that regulate neural progenitor cell (NPC) maintenance, proliferation, differentiation, adhesion, and polarity. These observations suggest that proper development of the ventricular system depends heavily on maintaining the NPCs that line the ventricles. However, the mechanisms through which disruptions in NPC maintenance contribute to ventricular obstruction and ultimately hydrocephalus remain poorly understood.

NPCs within the ventricular zone (VZ) of the developing brain not only serve as a proliferative reservoir for new neurons and glia but also form a layer of polarized epithelial cells lining the walls of the ventricular system. Thus, NPCs must carefully balance proliferation and differentiation, replenishing the progenitors to preserve the structural integrity of the ventricular system and subsequently give rise to the ependymal lining. A core pathway that coordinates this balance is Notch signaling. In mammals, Notch pathway activity is mediated by interactions between five ligands (Jagged 1 and 2; Delta-like 1, 3, and 4) and four receptors (Notch 1-4), which exhibit distinct spatial and temporal expression patterns throughout the central nervous system (CNS)^13–15^. Ligand engagement triggers γ-secretase-mediated cleavage of the Notch receptor, releasing the Notch intracellular domain (NICD), which translocates to the nucleus and binds to the transcriptional effector recombination signal binding protein for immunoglobulin kappa J region (RBPJ)^16,17^. Functioning as a transcriptional switch, this interaction displaces co-repressors and recruits co-activators to drive expression of downstream Notch targets, including the Hes family of basic helix-loop-helix (bHLH) transcription factors, notably *Hes1* and *Hes5*^18–21^. Within the CNS, Notch signaling plays a critical role in maintaining NPC populations and suppressing premature neurogenesis^22–27^. Notch signaling has also been shown to influence neural patterning by modulating a progenitor cell’s responsiveness to endogenous morphogens^28–31^. Lastly, beyond cell fate specification, Notch signaling has been shown to regulate mammalian cell adhesion and polarity^32–35^.

Previous studies in mice have demonstrated that elevated Notch signaling can lead to hydrocephalus through distinct pathogenic mechanisms. Loss of Rho family guanosine triphosphatase 3 (*Rnd3*) impairs ubiquitin-mediated degradation of NICD, thereby elevating Notch activity. In *Rnd3^-/-^*mice, excessive Notch signaling drives abnormal overgrowth of the ependymal epithelium, leading to physical obstruction of the cerebral aqueduct and ultimately obstructive hydrocephalus^36^. Similarly, loss of sorting nexin 27 (SNX27) increases γ-secretase activity, thereby elevating NICD production. However, the hydrocephalus in *Snx27^-/-^* mice results from defective ependymal cell differentiation and impaired ciliogenesis, leading to reduced ciliary beating and disrupted CSF flow^37,38^. In both studies, hydrocephalus arises as a downstream consequence of elevated Notch signaling, and pharmacological inhibition was sufficient to ameliorate the phenotype.

In this study, we demonstrate that preservation of Notch signaling also plays a critical role in the development of the ventricular system. We show that conditional loss of Notch signaling in the diencephalon results in premature neuronal differentiation and depletion of NPCs from the VZ. This also leads to deficits in the formation of ependymal cells lining the ventricles, resulting in collapse of its walls and obstruction of CSF flow. In a parallel set of experiments, we show that pharmacological inhibition of Notch signaling disrupts neural rosette formation in a cortical spheroid system, recapitulating aspects of neuroepithelial defects observed in vivo. Collectively, our findings demonstrate that proper maintenance of NPCs within the VZ is critical for preserving the structural integrity of the ventricular system and maintaining an unobstructed path for CSF flow. Altogether, the results show the importance of balanced Notch signaling as perturbations in either direction can compromise ventricular architecture and ultimately lead to hydrocephalus.

## RESULTS

### Conditional loss of *Rbpj* reveals region-specific dependence on Notch signaling

RBPJ is an essential transcriptional effector of canonical Notch signaling, making it an ideal target for pathway inactivation. Complete loss of *Rbpj* results in delayed growth and early embryonic lethality (e8.5-e10.5)^39^, preventing analysis of its roles during later stages of brain development. To circumvent this limitation, we selectively disrupted Notch signaling in defined NPC populations in the developing forebrain and midbrain by generating *Rbpj* conditional knockout (*Rbpj^CKO^*) mice using an *Olig2^Cre^* driver^40^. These animals were crossed to a Cre-inducible *R26R^GFP/GFP^* reporter^41^ to label recombined cells and to an *Rbpj* floxed allele (*Rbpj^flox/flox^*)^42^ to enable conditional gene deletion. As designed, at age e12.5, GFP expression robustly overlapped with SOX2+ and OLIG2+ NPCs in control embryos (**Figure 1A-K**), confirming *Olig2^Cre^*-mediated recombination along various regions of the developing ventricular system.

**Figure 1:**
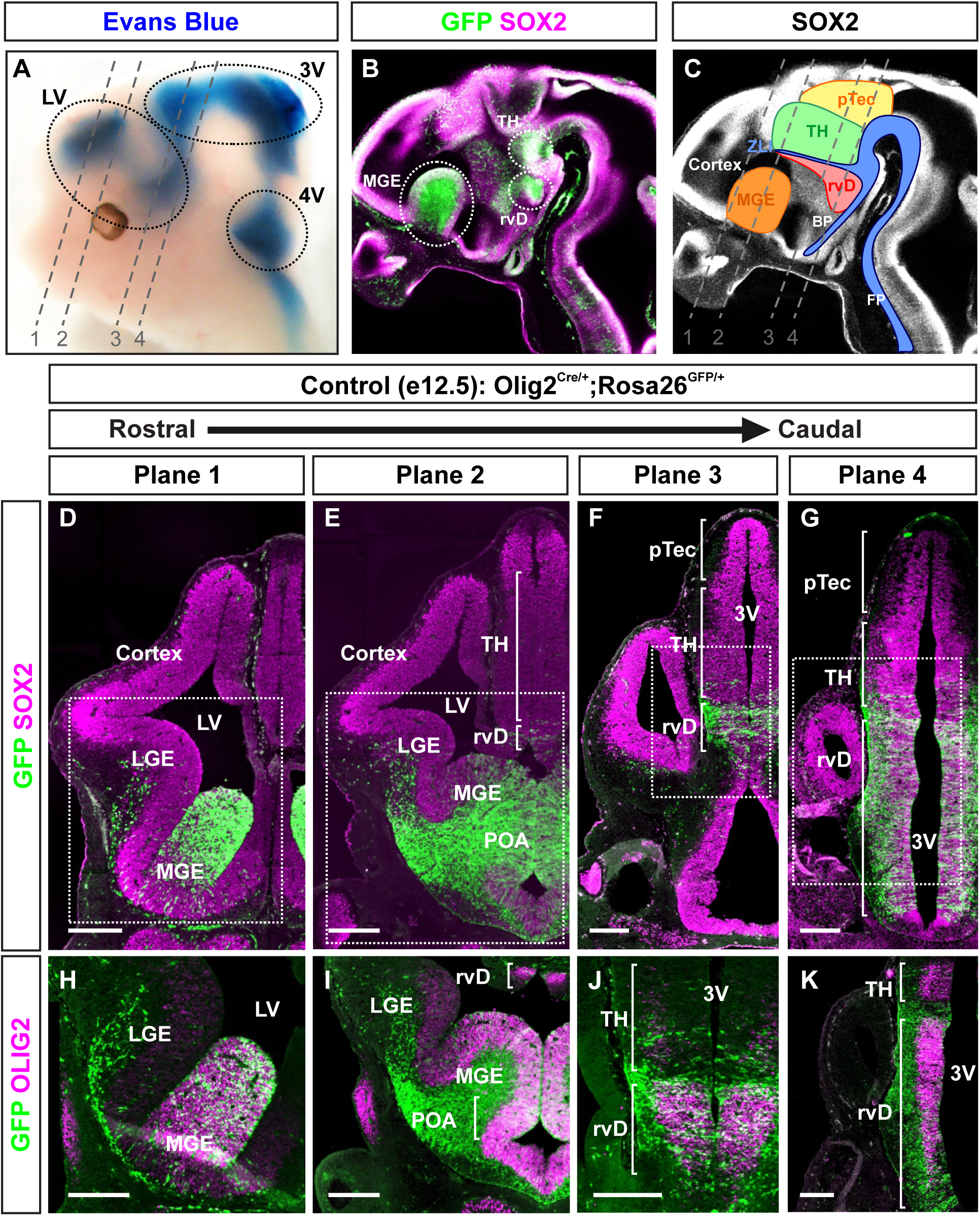
Spatial analysis of *Olig2^Cre^* mediated recombination in the developing brain. **(A-C)** Lateral view of the head of an e12.5 control mouse embryo (*Olig2^Cre/+^*; *R26^GFP/+^*). Dashed grey lines (labeled 1-4) indicate the planes of coronal sections captured in D-K. **(A)** Evans blue dye injected into the lateral ventricles (LV) to visualize the ventricular system, including the third ventricle (3V) and fourth ventricle (4V). **(B)** Whole-mount immunofluorescence (IF) for SOX2 and GFP. Dashed white circles identify major GFP+ populations denoting regions of Cre-mediated recombination within the telencephalon and diencephalon. **(C)** Schematic illustrating the location of the basal plate (BP), floor plate (FP), medial ganglionic eminence (MGE), preoptic area (POA), pretectum (pTec), rostral/ventral diencephalon (rvD), thalamus (TH), and zona limitans intrathalamica (ZLI). **(D-G)** Coronal sections through planes 1-4 stained for SOX2 and GFP. Dashed white boxes indicate the regions shown at higher magnification in H-K. Scale bars: 200 µm. **(H-K)** Higher magnification IF images stained for OLIG2 and GFP. Scale bars: 200 µm.

To verify a conditional loss of Notch signaling activity, we first looked at the expression of the downstream target *Hes5* and assessed its effects on NPC maintenance in GFP+ areas (**Figure 2**). Unexpectedly, *Rbpj* deletion produced region-specific effects. In the telencephalon, a loss of *Rbpj* in the medial ganglionic eminence (MGE) (Plane 1) did not significantly alter *Hes5* expression or SOX2+ NPC abundance (**Figures 2B-C, 2F-G, 2J-K**). In contrast, just caudal to this region (Plane 2), *Rbpj* loss within the preoptic area (POA) extinguished *Hes5* expression, accompanied by modest thinning of SOX2+ NPCs (**Figures 2D-E, 2H-I, 2L-M**). Lastly, in regions of the diencephalon (Plane 4), deletion of *Rbpj* resulted in a more severe phenotype characterized *Hes5* loss and marked depletion of SOX2+ NPCs (**Figures 2N-S**). Collectively, these findings demonstrate that *Olig2^Cre^*-mediated deletion of *Rbpj* successfully inhibits canonical Notch signaling in discrete regions of the developing brain. However, our analysis also revealed unexpected regional differences in pathway dependence, with *Rbpj* loss having stronger effects on NPCs within the POA and rostral/ventral diencephalon (rvD) compared to the MGE.

**Figure 2:**
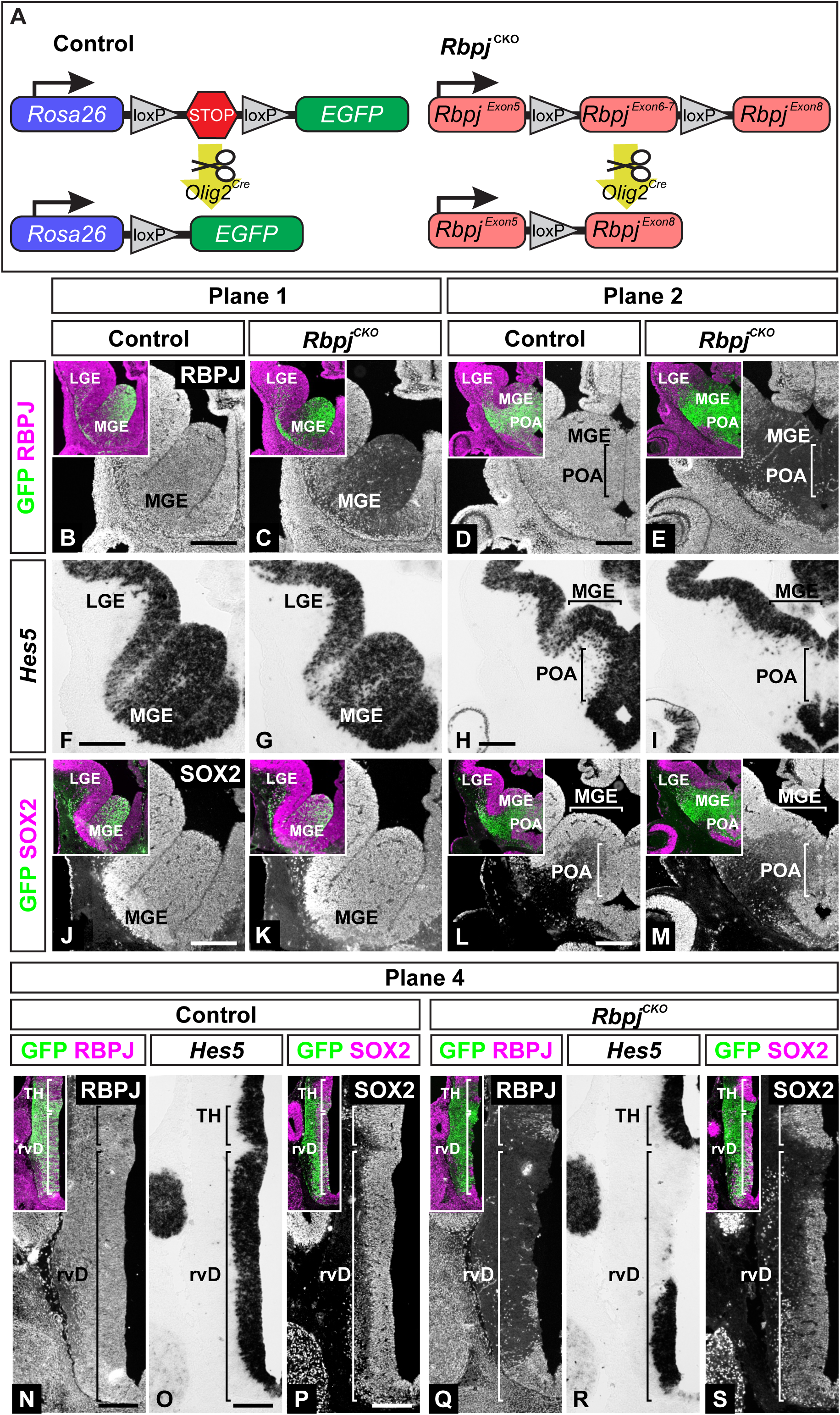
Loss of *Rbpj* results in region-specific changes in Notch signaling activity and NPC maintenance. **(A)** Schematic of *Olig2^Cre^*-mediated mouse manipulations. (Left) In control mice (*Olig2^Cre/+^*; *R26^GFP/+^*), a *Rosa26^GFP^*reporter identifies cells that undergo Cre-mediated recombination. (Right) In *Rbpj^CKO^* mice, *Olig2^Cre^*-mediated recombination removes exons 6 and 7 of *Rbpj*, resulting in loss of functional RBPJ. **(B-S)** Coronal sections of the telencephalon and diencephalon of e12.5 control and *Rbpj^CKO^* brains corresponding to planes 1, 2, and 4 defined in Figure 1. Immunofluorescence (IF) images of RBPJ, SOX2, and GFP. Insets highlight the GFP+ regions of Cre-mediated recombination. *Hes5* expression is assessed by in situ hybridization (ISH). Lateral ganglionic eminence (LGE), medial ganglionic eminence (MGE), preoptic area (POA), rostral/ventral diencephalon (rvD), and thalamus (TH). Scale bars: 200 µm.

### Depletion of NPCs disrupts the VZ in *Rbpj^CKO^* mice

To better understand how loss of Notch signaling affects the developing ventricular system, we focused on the diencephalon surrounding the third ventricle (Plane 4) (**Supplementary Figure S1**). In this region, e12.5 *Rbpj^CKO^* mutants consistently displayed a complete depletion of SOX2+ NPCs within the VZ, accompanied by a corresponding increase in TUJ1+ neurons (**Figures 3A-H**). Consistent with this loss of progenitors, there was also a reduction in Jagged1 (JAG1), a Notch ligand expressed by NPCs in this region^15^ (**Figures 3I-J**). As NPCs are highly polarized epithelial cells maintained by junctional adhesion complexes^43–46^, we next assessed how their premature differentiation into neurons affected the composition of the VZ. Rather than undergoing orderly delamination and migration away from the ventricular surface, newly differentiated neurons disrupted continuity in the neuroepithelium, creating a prominent gap in progenitors lining the ventricles. Consistent with a loss of NPCs, acetylated α-tubulin (*α*TUB) staining within this gap revealed a loss of columnar cell morphology (**Figures 3K-L**), and ARL13B staining showed a loss of apically localized primary cilia (**Figures 3M-N**). In addition, the polarity marker atypical protein kinase C (aPKC) (**Figures 3O-P**) and the adherens junction proteins N-cadherin (NCAD) and β-catenin (βCAT) (**Figures 3Q-T**) were all reduced at the apical surface. Importantly, this loss of neuroepithelial continuity occurred after the VZ had already formed. *Olig^Cre^*-mediated deletion of *Rbpj* occurs after the VZ has been established. At e10.5, the VZ is intact, with SOX2+ and OLIG2+ NPCs still present and with no overt defects in polarity or cell-to-cell adhesion contacts (**Supplementary Figure S2**). Collectively, these findings show that sustained Notch signaling is required in this region of the brain to maintain the NPC population and preserve neuroepithelial integrity. When Notch signaling is lost, premature neuronal differentiation depletes progenitors, disrupts VZ continuity, and these defects persist for the remainder of brain development (**Supplementary Figure S3**).

**Figure 3:**
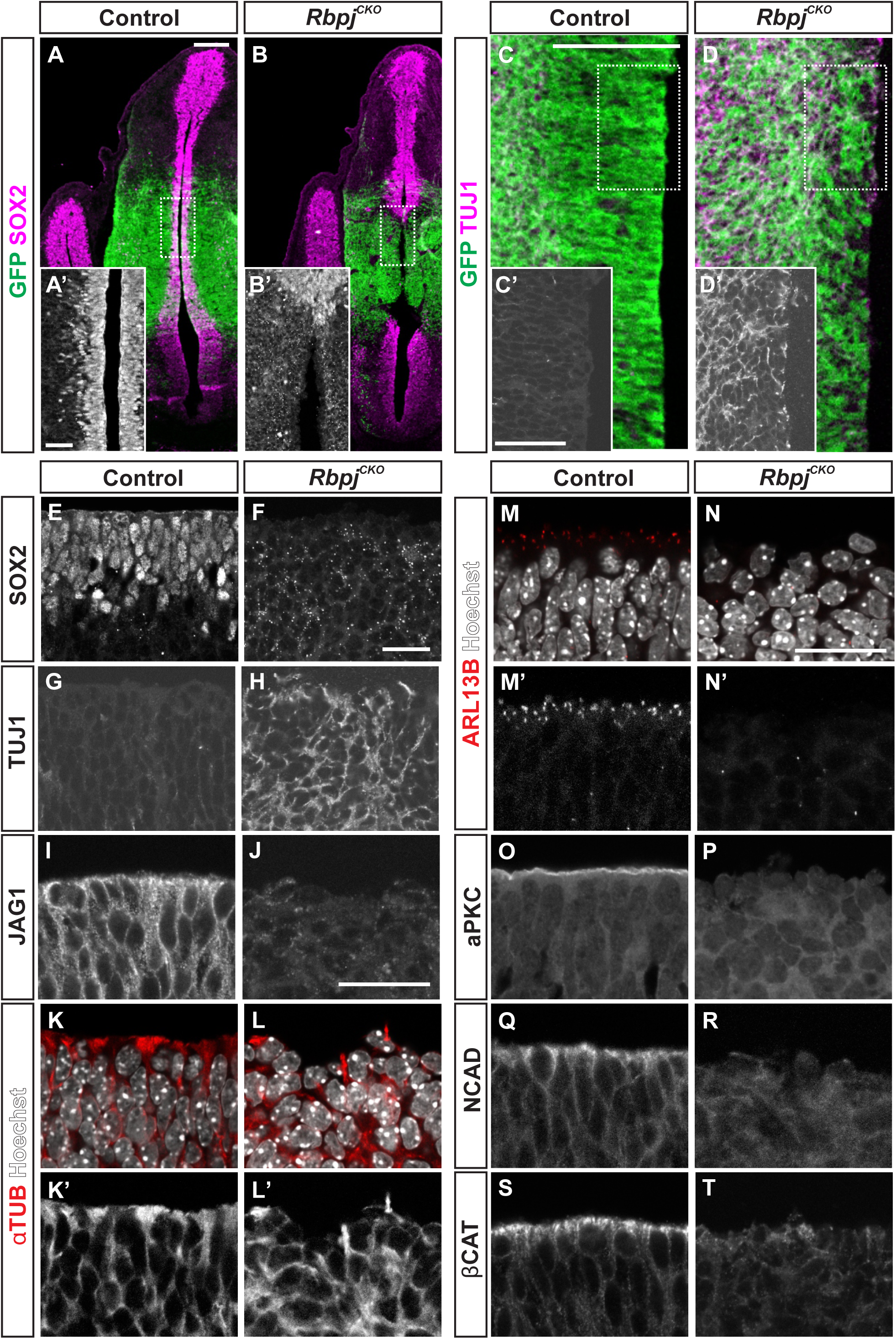
Premature neuronal differentiation disrupts ventricular zone integrity in the developing diencephalon. **(A-D)** Low-magnification coronal sections through the rostral/ventral diencephalon (rvD) of e12.5 control and *Rbpj^CKO^* embryos. (A-B) Immunofluorescence (IF) of SOX2 and GFP. Enlarged images (A’-B’) correspond to the regions within the dashed white boxes. (C-D) IF of TUJ1 and GFP. Enlarged images (C’-D’) correspond to the regions within the dashed white boxes. The apical surface is oriented to the right. Scale bars: 200 µm (low mag) and 50 µm (high mag). **(E-T)** High-magnification analysis of cell identity, polarity, and adhesion within the VZ. (E-F) SOX2+ NPCs. (G-H) TUJ1+ neurons. (I-J) Jagged1 (JAG1), a Notch ligand expressed by NPCs in this region. (K-L) Acetylated α-tubulin (αTUB), which labels cytoplasmic microtubules and outlines cell morphology. (M-N) ARL13B+ primary cilia concentrated at the apical surface of NPCs and extending into the ventricle. (O-P) aPKC, an apically localized polarity marker and component of the Par3/Par6/aPKC complex. (Q-R) NCAD and (S-T) βCAT, apically enriched adherens junction proteins. In all higher-magnification images (E-T), the apical surface is oriented upward. Scale bars: 25 µm.

### Loss of NPCs in embryonic *Rbpj^CKO^* mice results in postnatal ependymal defects

Ependymal cells form a single ciliated epithelial layer that lines the ventricular system and plays a major role in regulating CSF movement and homeostasis^47–49^. These cells are derived from NPCs and mature postnatally into multiciliated cells^50^. Given the severe disruption of the SOX2+ NPCs in the embryonic *Rbpj^CKO^*brains (**Figures 4A-C**), we asked how these early defects impacted the ependymal layer in the adult brain. In postnatal day 60 (P60) control brains, the ependymal layer lining the third ventricle was intact and clearly identified by the expression of the intermediate filament marker vimentin (VIM)^51^ (**Figure 4D**). In contrast, the ventricular lining in *Rbpj^CKO^* mutants appeared disorganized and exhibited few VIM+ cells (**Figure 4F**). Consistent with this finding, mutant brains also showed greatly reduced expression of the ependymal cell marker S100 calcium-binding protein B (S100B)^52^ (**Figures 4E, 4G**). Together, these findings indicate that loss of Notch signaling during embryonic development depletes ependymal cells at postnatal stages (**Figure 4H)**.

**Figure 4:**
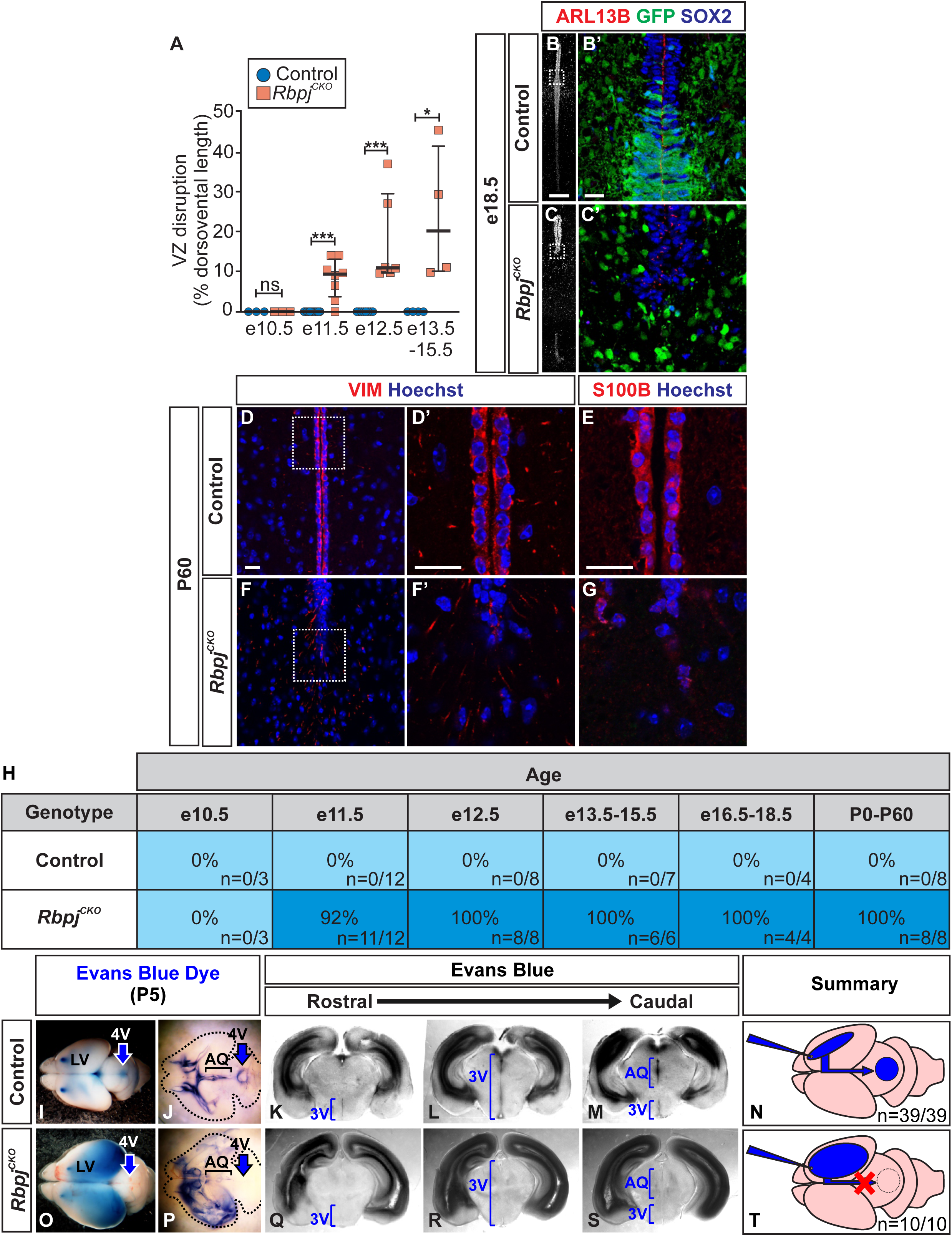
Neuroepithelial defects in *Rbpj^CKO^* mice persist into adulthood and obstruct CSF flow. **(A-F)** Coronal sections through the diencephalon of e18.5 and P60 control and *Rbpj^CKO^* brains. (A and D) Immunofluorescence (IF) of SOX2+ NPCs, ARL13B+ primary cilia, and GFP. Enlarged images (A’ and D’) correspond to the regions within the dashed white boxes. Scale bars: 200 µm (low mag) and 20 µm (high mag). (B-C and E-F) IF of VIM+ and S100B+ ependymal cells lining the third ventricle. Enlarged images (B’, C, E’, F) correspond to the regions within the dashed white boxes. Scale bars: 20 µm. **(G-R)** Evans blue dye injected into the lateral ventricles of P5 control and *Rbpj^CKO^* brains to assess CSF flow through the ventricular system. (G, M) External views of dye-filled brains before clearing. (H, N) Dye-filled brains after clearing. The black dashed line outlines the brain. (I-K, O-Q) Serial coronal vibratome sections of control and *Rbpj^CKO^* dye-filled brains. Blue brackets identify the location of the ventricular system. (L and R) Summary of the Evans Blue assay and an illustration predicting the location of the ventricular obstruction. Control (n = 39) and *Rbpj^CKO^* (n = 10). Lateral ventricles (LV), third ventricle (3V), cerebral aqueduct (AQ), and fourth ventricle (4V). **(S)** Summary of control and *Rbpj^CKO^* brains examined across embryonic and postnatal ages for premature neuronal differentiation and loss of NPC continuity along the VZ.

### Ependymal defects in *Rbpj^CKO^* mice obstruct the ventricular system

The observed loss of the ependymal layer coincided with apparent closures within the third ventricle as surrounding tissue extended across the space normally occupied by the ventricular lumen (**Figures 4D-G**). To confirm that this disruption to the ventricular system indeed obstructed CSF flow, we injected Evans Blue dye into the lateral ventricles of extracted neonatal P5 brains to visualize CSF flow. In control brains, the dye filled the lateral ventricles and readily propagated through the ventricular system (**Figures 4I-J**). In contrast, the dye accumulated within the enlarged lateral ventricles of *Rbpj^CKO^*brains and failed to progress further, indicating a physical obstruction (**Figures 4O-P**). All mutant brains analyzed using the Evans Blue assay displayed obstructed ventricular systems and enlarged lateral ventricles (Control n = 39, *Rbpj^CKO^*n = 10). To locate the site of the obstruction, brains injected with dye were fixed and sectioned (**Figures 4K-N**). In *Rbpj^CKO^* brains, the dye was absent beyond the lateral ventricles, indicating the blockage occurred within the cerebral aqueduct or third ventricle (**Figures 4Q-T**). Thorough analysis of *Rbpj^CKO^* brains across embryonic and postnatal ages revealed that some disruption of the VZ occurred as early as e11.5, and the consequences to the ependymal cell layer consistently persisted into adulthood. Collectively, these findings suggest that loss of Notch signaling in *Rbpj^CKO^* mice causes an early loss of NPCs that compromises the integrity of the ventricular system, eventually leading to closure of the ventricular lumen and obstructed CSF flow.

### *Rbpj^CKO^* mice develop hydrocephalus

The ventricular obstruction observed in neonatal *Rbpj^CKO^*mice can result in hydrocephalus. While P5 *Rbpj^CKO^* mice were approximately 15% smaller than controls, they displayed a slightly enlarged head (**Figures 5A-C**). As the animals aged, the phenotypes became more pronounced. By weaning (P20-P26), the *Rbpj^CKO^*mice weighed ∼55% less than controls and ∼45% exhibited the characteristic cranial doming associated with severe hydrocephalus (**Figures 5D-I**). Severely affected *Rbpj^CKO^* animals developed impaired locomotor and feeding behaviors. Interestingly, the absence of cranial doming did not indicate an absence of ventricular pathology. A closer look revealed that *Rbpj^CKO^* mice without domed heads still had modestly enlarged ventricles, consistent with the high penetrance of VZ defects and CSF obstruction observed in mutant animals. These findings indicate that although Notch signaling loss causes highly penetrant defects in ventricular development in *Rbpj^CKO^*mice, the resulting hydrocephalus varies in severity.

**Figure 5:**
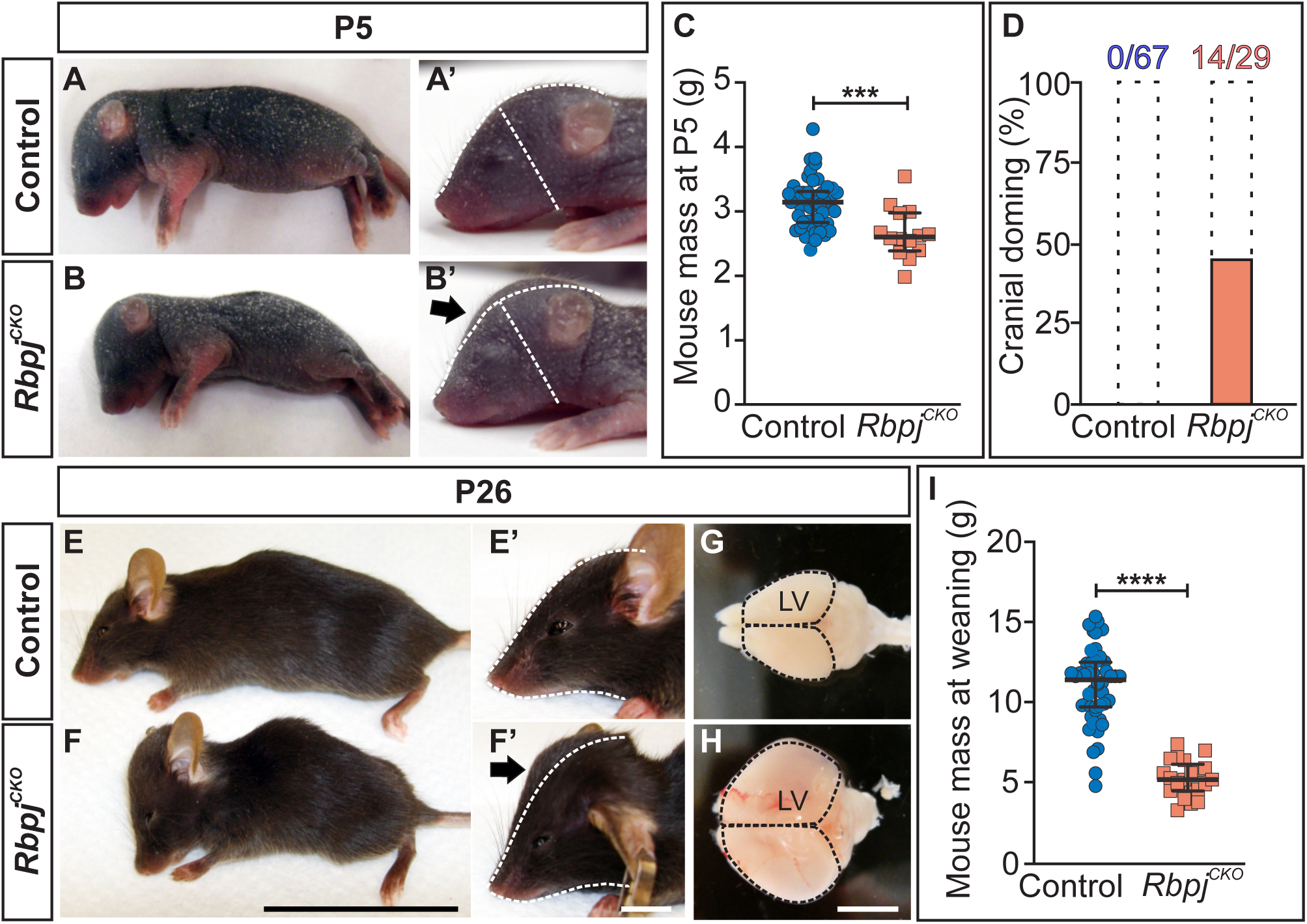
*Rbpj^CKO^* mice develop hydrocephalus. **(A-B)** Neonatal (P5) control and *Rbpj^CKO^* mice. (A’-B’) Enlarged images of the heads. Dashed white lines are the outline of the control skull, superimposed on the *Rbpj^CKO^* skull to highlight cranial doming (black arrow). **(C)** Scatter dot plot of P5 body mass. Bars represent the median with interquartile range. Median control: 3.14 g (n = 46); median *Rbpj^CKO^*: 2.61 g (n = 14). Mann-Whitney test, ***p = 0.0001. **(D)** Penetrance of cranial doming observed at weaning (P20-26) in control (n = 67) and *Rbpj^CKO^* (n = 29) mice. **(E-H)** Recently weaned (P20-26) control and *Rbpj^CKO^* mice. (E’-F’) Enlarged images of the heads. Dashed white lines are the outline of the control skull, superimposed on the *Rbpj^CKO^* skull to highlight cranial doming (black arrow). Scale bars: 5 cm (low mag) and 1cm (high mag). (G-H) Brains collected from control and *Rbpj^CKO^* mutants. The dashed black lines outline the brain to better illustrate the enlarged lateral ventricles (LV). Scale bar: 1 cm. **(I)** Scatter dot plot of weaned (P20-26) body mass. Bars represent the median with interquartile range. Median control: 11.4 g (n = 48); median *Rbpj^CKO^* = 5.2 g (n = 21). Mann-Whitney test, ****p < 0.0001.

### Notch inhibition disrupts neural rosette organization in cortical spheroids

Given the growing number of candidate mutations associated with congenital hydrocephalus, experimental models are needed to assess their effects on brain development. We therefore asked if we could reproduce phenotypes observed in *Rbpj^CKO^* mice in a cortical spheroid system. To approximate the timing of Notch inhibition in the mouse model, we began using a pharmacological approach. For these experiments, mESCs were plated into non-adherent wells and differentiated into cortical spheroids. On day 6 of differentiation, the cortical spheroids were treated with either a vehicle control (DMSO) or the γ-secretase inhibitor Compound E (CompE) and allowed to grow for an additional 4 days (**Figure 6A**). CompE addition strongly reduced the expression of the Notch target *Hes5* (**Figure 6B**). Control spheroids readily formed neural rosettes, radial arrangements of NPCs that self-organize around a central apical ring and model aspects of neuroepithelial development. Rosettes were identified by the presence of a central ring of the tight junction protein zona occludens 1 (ZO1) (**Figures 6C-E**). Quantification of these ZO1+ structures revealed significantly fewer rosettes following CompE treatment (**Figure 6C**). We also examined NCAD and the organization of ARL13B+ primary cilia to assess the ability of CompE-treated spheroids to form neural rosettes. Consistent with the ZO1 analysis, expression of these proteins was largely absent in CompE-treated spheroids (**Figures 6E-J**). Collectively, these results show that continued Notch signaling is needed to preserve neuroepithelial architecture in cortical spheroids and suggest that cortical spheroid cultures can serve as a convenient platform for initially assessing how candidate mutations associated with hydrocephalus disrupt early brain development.

**Figure 6:**
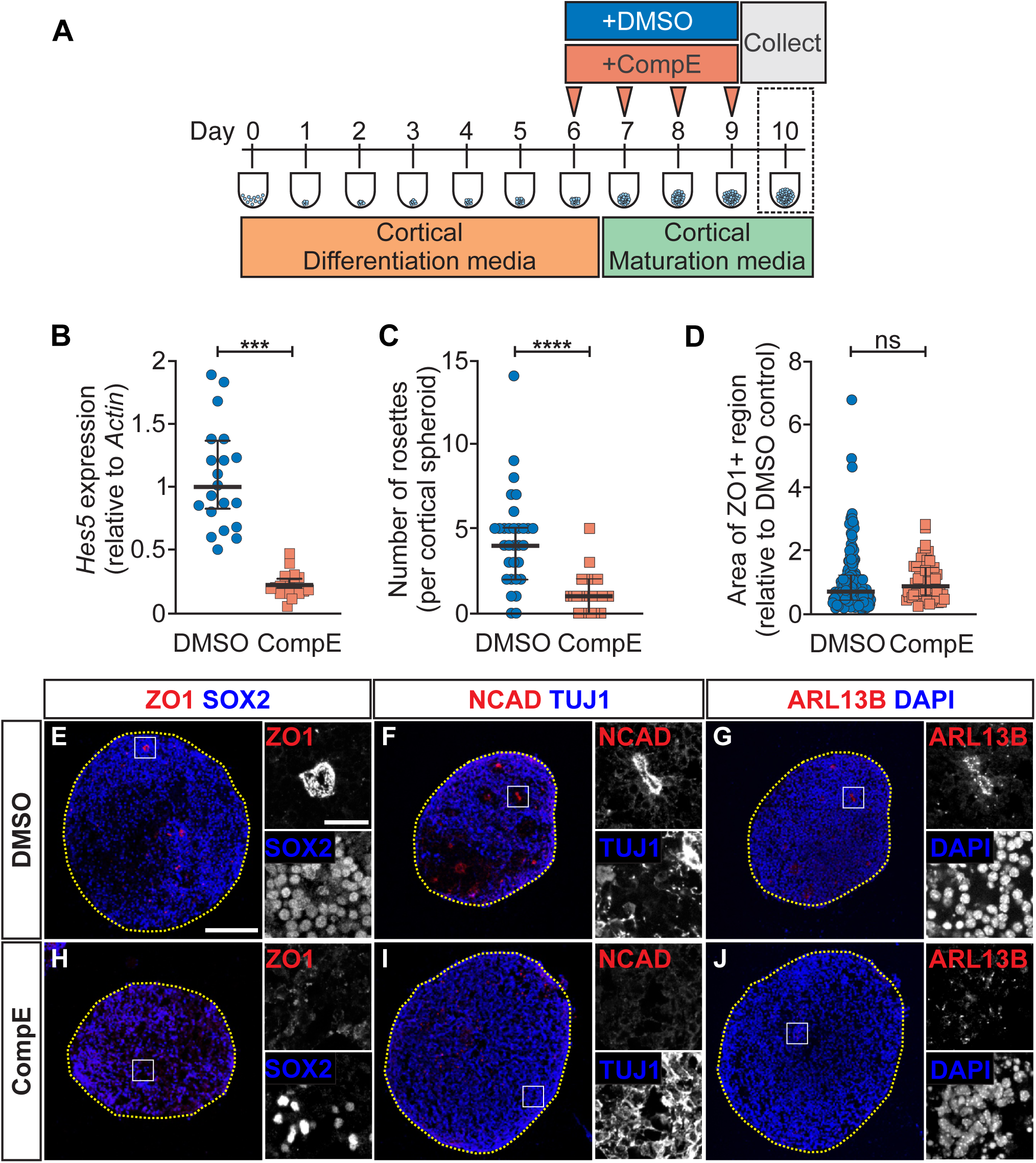
Pharmacological inhibition of Notch signaling activity disrupts neural rosette organization in cortical spheroids. **(A)** Schematic of the cortical spheroid differentiation protocol. mESCs are differentiated into cortical spheroids for 10 days. On day 6, spheroids are treated with either vehicle control (DMSO) or the γ-secretase inhibitor Compound E (CompE, 10 nM) and collected on day 10 for analysis. **(B)** *Hes5* expression in cortical spheroids treated with either DMSO or CompE and normalized to *Actin*. Each data point represents a technical replicate from 7 independent experiments. Bars represent the median with interquartile range. Median DMSO: 1.00 (n = 7); median CompE: 0.22 (n = 7). Mann-Whitney test, ***p = 0.0006. **(C)** Quantification of ZO1+ neural rosettes. Each data point represents the number of rosettes counted within a single spheroid section collected across three independent experiments. Bars represent the median with interquartile range. Median DMSO: 4 rosettes (n = 30), median CompE: 1 rosette (n = 29). Mann-Whitney test, ****p < 0.0001. **(D)** Quantification of individual ZO1+ region areas. Each data point represents the area of a single ZO1+ region across three independent experiments. Values from each experiment are normalized to the mean ZO1+ area of the corresponding DMSO controls to account for experiment-to-experiment variability in spheroid size. Bars represent the median with interquartile range. Median DMSO: 0.69 (n = 188); median CompE = 0.87 (n = 54). Mann-Whitney test, ns, p = 0.0509. **(E-J)** Immunofluorescence (IF) images of cortical spheroids treated with DMSO or CompE. Spheroids are stained for (E, H) ZO1 and SOX2, (F, I) NCAD and TUJ, and (G, J) ARL13B and DAPI. Yellow dashed lines outline the spheroids. Dashed white boxes indicate the regions shown at higher magnification to the right. Scale bars: 200 µm (low mag) and 20 µm (high mag).

We next examined the consequences of a complete loss of *Rbpj* in this cortical spheroid platform. Using CRISPR-Cas9 editing, we generated *Rbpj^KO^*mESCs and confirmed the loss of RBPJ protein by Western blot and reduced Notch signaling by *Hes5* expression (**Supplementary Figures S4A-C**). We then differentiated these *Rbpj^KO^* mESCs into cortical spheroids and compared them to spheroids generated from mESCs that received a non-targeting control. Control cells formed well-organized neural rosettes with a central ring of ZO1. Surprisingly, unlike CompE-treated spheroids, which formed fewer rosettes, *Rbpj^KO^* spheroids had clusters of expanded ZO1 expression (**Supplementary Figures 4D-G**). Thus, although both approaches disrupted Notch signaling and altered rosette formation, they did so in distinct ways. Collectively, these findings show that while Notch signaling is important for neural rosette organization in cortical spheroids, different phenotypes can arise depending on when and how the pathway is disrupted.

## DISCUSSION

Hydrocephalus is one of the most common congenital brain malformations worldwide^53^ and remains a leading cause of pediatric neurosurgery. Although early surgical intervention can restore CSF circulation, many patients continue to experience persistent neurological complications^54–57^, suggesting that some of the underlying pathology may extend beyond elevated intracranial pressure. Recent genetic studies support this idea^7–12^, with many candidate mutations predicted to disrupt NPC maintenance, proliferation, adhesion, and differentiation. This raises an important question: how does disrupting NPC biology contribute to congenital hydrocephalus? NPCs not only serve as a source of new neurons and glia for the developing brain, but they also form a polarized neuroepithelial layer lining the developing ventricular system and ependymal cell layer in the adult brain. Thus, their differentiation presents a unique architectural challenge. Differentiation, delamination, and migration of progeny must all occur without compromising the continuity of NPCs that line the ventricular wall. Our studies show that premature differentiation can deplete NPCs and create gaps in the VZ, disrupting the integrity of the ventricular system and ultimately obstructing CSF flow.

Notch signaling is a central regulator of NPC maintenance^22,27^. In this study, we show that conditional loss of *Rbpj* results in premature neurogenesis and depletion of NPCs within the VZ. Defects in the neuroepithelial layer during embryogenesis have a lasting impact on the postnatal ependymal lining, ultimately contributing to obstructive hydrocephalus (**Figure 7**). Thus, our findings link the established role of Notch signaling in NPC maintenance to a broader requirement for preserving the architecture of the developing ventricular system. Interestingly, the consequences of *Rbpj* loss were not uniform across the developing brain. While the rostral/ventral diencephalon exhibited a striking loss of SOX2+ NPCs, progenitors within the medial ganglionic eminence (MGE) were preserved, and no disruptions in the ventricular system were detected. Similar regional differences have been observed in the developing spinal cord, where *Rbpj* deletion results in progenitor loss and disruption of the VZ in intermediate domains, while continuity remained intact in more ventral domains^29,58^. These findings suggest that the requirement of Notch signaling in maintaining progenitor populations varies across the developing CNS. One possibility is that progenitor maintenance in more resilient regions may be supported by Notch-independent pathways. For example, Hedgehog and FGF signaling can stimulate the expression of *Hes*/*Hey* transcription factors and may provide compensatory signals that help maintain NPCs following *Rbpj* loss^59–61^.

**Figure 7:**
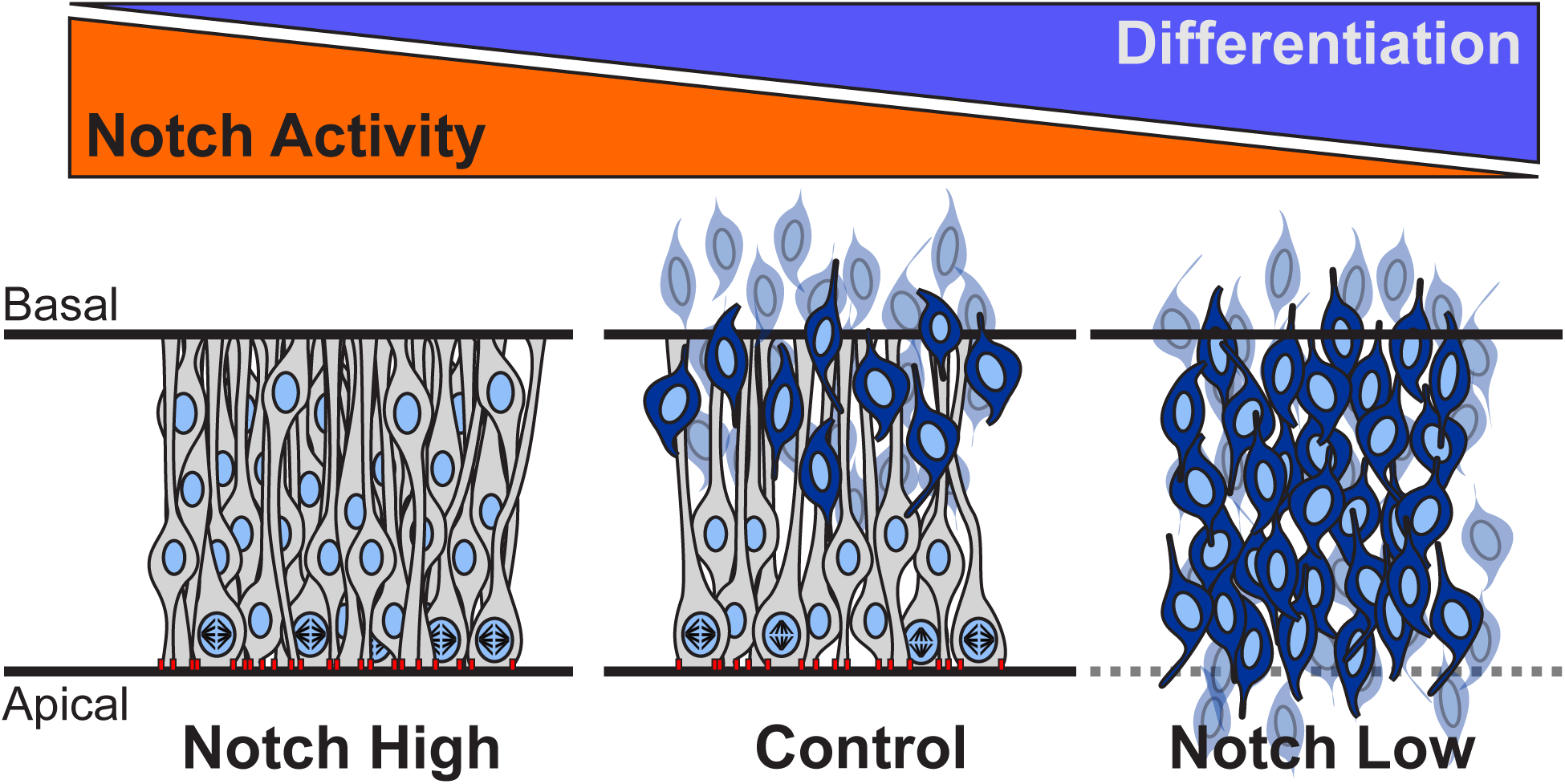
Model of the relationship between Notch signaling activity and the maintenance of apical cell adhesion and cell polarity of NPCs within the diencephalon. Schematic illustrating the effects of elevated, normal, and reduced Notch signaling activity on NPC maintenance and the organization of the VZ. With elevated Notch signaling activity (Notch High), cells are retained as NPCs, and neuronal differentiation is reduced. At normal levels of Notch signaling activity (Control), NPC maintenance is balanced with neuronal differentiation. In the absence of Notch signaling (Notch Low), premature neuronal differentiation depletes NPCs and disrupts the VZ, resulting in a loss of apically localized cell adhesion and polarity proteins.

Our findings also highlight the importance of balancing NPC maintenance and differentiation for ventricular system development, as both elevated and reduced Notch signaling activity can result in hydrocephalus. Previous studies have shown that hyperactivation of Notch signaling, including through loss of the Notch antagonist *Rnd3*, leads to excessive NPC proliferation, aqueductal stenosis, and consequently obstructive hydrocephalus^36^. Consistent with these findings, we observed that hyperactive Notch signaling through conditional expression of NICD in *Olig2*-expressing cells (*Rosa^NICD^*) led to a similar expansion of SOX2+ NPCs into the ventricular space (**Supplementary Figures S5**). In contrast, loss of *Rbpj* had the opposite effect, causing premature neuronal differentiation and NPC depletion. Despite their opposing effects, elevated and reduced Notch signaling activity both alter the NPC landscape and thus the abundance of progenitors lining the ventricular system. Collectively, research from our lab and others suggests that ventricular system development is highly sensitive to Notch signaling dosage. Shifting Notch signaling activity in either direction can disrupt ventricular wall architecture and consequently obstruct CSF flow.

Although approximately 95% of congenital hydrocephalus cases are sporadic with no known causal variant^62^, large-scale genomic studies have implicated hundreds of candidate mutations^7,8,10–12,63,64^. Many of these mutations are predicted to disrupt core developmental signaling pathways, including Notch, Hedgehog, EGF, FGF, and Wnt. These pathways regulate many processes, including NPC cell proliferation, adhesion, polarity, and differentiation. A major challenge, however, is validating these candidate mutations, determining the processes they disrupt, and ultimately how they contribute to hydrocephalus. To begin addressing this challenge, we used cortical spheroids to determine whether perturbing Notch signaling could reproduce aspects of the neuroepithelial defects we observed in vivo. Pharmacological inhibition of Notch signaling mid-differentiation reduced neural rosette formation and disrupted neuroepithelial organization, closely resembling the phenotypes observed in *Rbpj^CKO^* mice. Interestingly, constitutive loss of *Rbpj* produced a different phenotype. Rather than reducing rosette numbers, *Rbpj^KO^* spheroids developed enlarged and disorganized ZO1+ regions. An important difference between these experiments is the timing of Notch inhibition. These findings raise the possibility that the consequences of Notch signaling inhibition depend on the developmental stage at which signaling is disrupted. These findings provide a starting point for using cortical spheroids as a tractable model to evaluate how hydrocephalus-associated mutations affect early brain development and to prioritize candidates for further in vivo investigation. The platform might have additional value in facilitating screens for compounds and interventions that could preserve neuroepithelial architecture across development.

## ACKNOWLEDGEMENTS

The authors thank Dr. Wai Pang Chan and the UW Biology Imaging Facility for their important contributions to this work.

## AUTHOR CONTRIBUTIONS

Conceptualization: L.G., L.N., T.M., D.M., D.L., C.A.P., B.G.N., J.H.K.

Methodology: L.G., L.N., T.M., C.E.D., C.A.P., B.G.N., J.H.K.

Software:

Validation: D.L., D.M., G.S., A.I.L.O

Formal analysis: L.N., T.M., D.L., D.M., G.S., A.I.L.O, J.H.K.

Investigation: L.G., L.N., T.M., D.L., D.M., C.E.D., G.S., A.I.L.O, C.A.P., J.H.K.

Resources: C.A.P., B.G.N., J.H.K.

Data curation:

Writing (original draft preparation): L.G., L.N., C.A.P., B.G.N, J.H.K.

Writing (review and editing): C.A.P., B.G.N, J.H.K.

Visualization: L.G., L.N., T.M., J.H.K.

Supervision: C.A.P., B.G.N., J.H.K.

Project administration: C.A.P., B.G.N., J.H.K.

Funding acquisition: C.A.P., B.G.N., J.H.K.

## DATA AVAILABILITY

All relevant data are available in the article and its supplementary information.

---

**Supplementary Figure S1.**
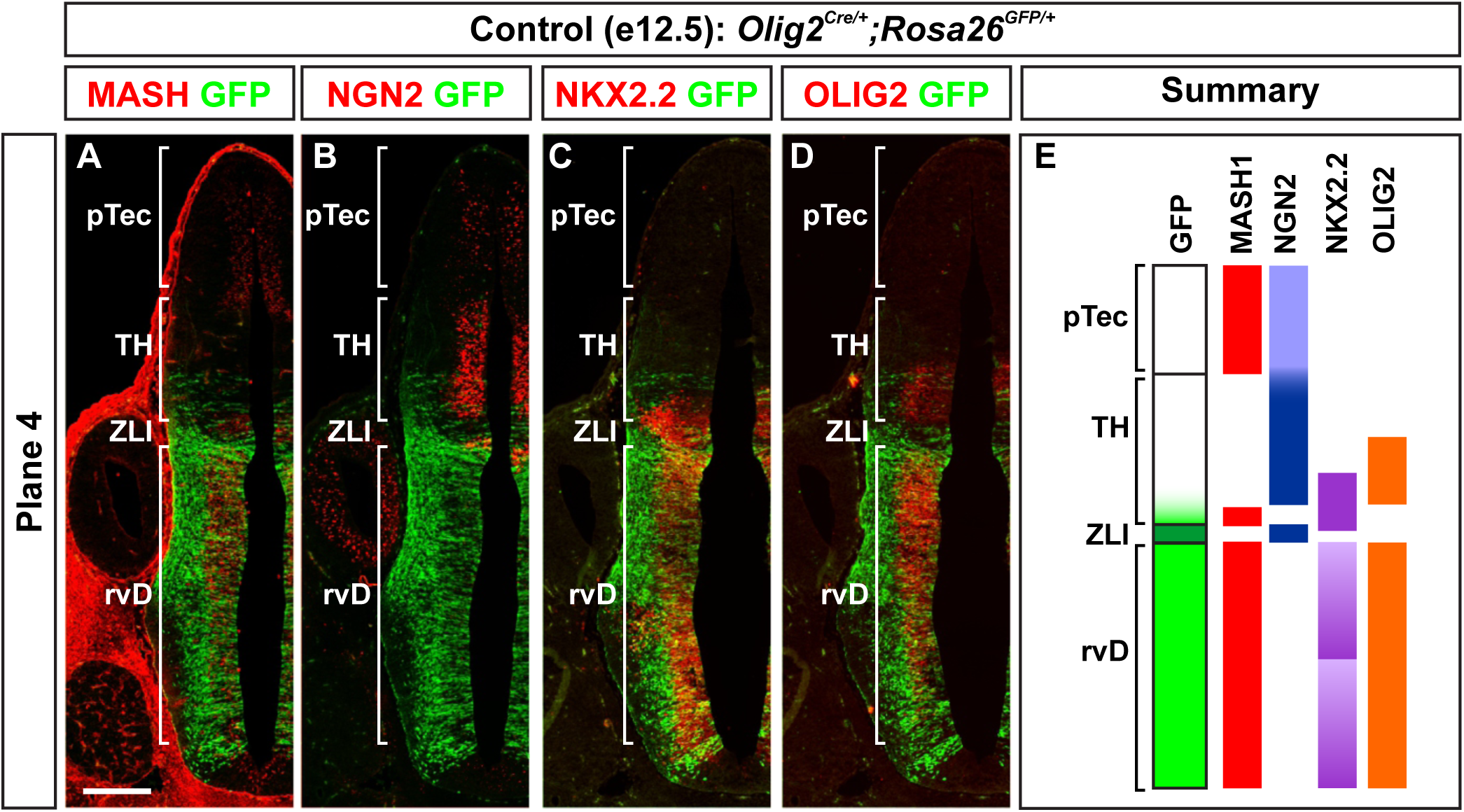
Detailed analysis of Olig2^Cre^ mediated recombination in the developing diencephalon. **(A-D)** Coronal sections of e12.5 control brains corresponding to plane 4 as defined in Figure 1. Immunofluorescence (IF) for GFP and the transcription factors MASH1, NGN2, NKX2.2, and OLIG2. Combinatorial expression of these transcription factors can identify distinct progenitor populations within the developing diencephalon^53,54^. **(E)** Summary of where *Olig2^Cre^*-mediated recombination occurs. Pretectum (pTec), rostral/ventral diencephalon (rvD), thalamus (TH), and zona limitans intrathalamica (ZLI). Scale bars: 200 µm.

**Supplementary Figure S2.**
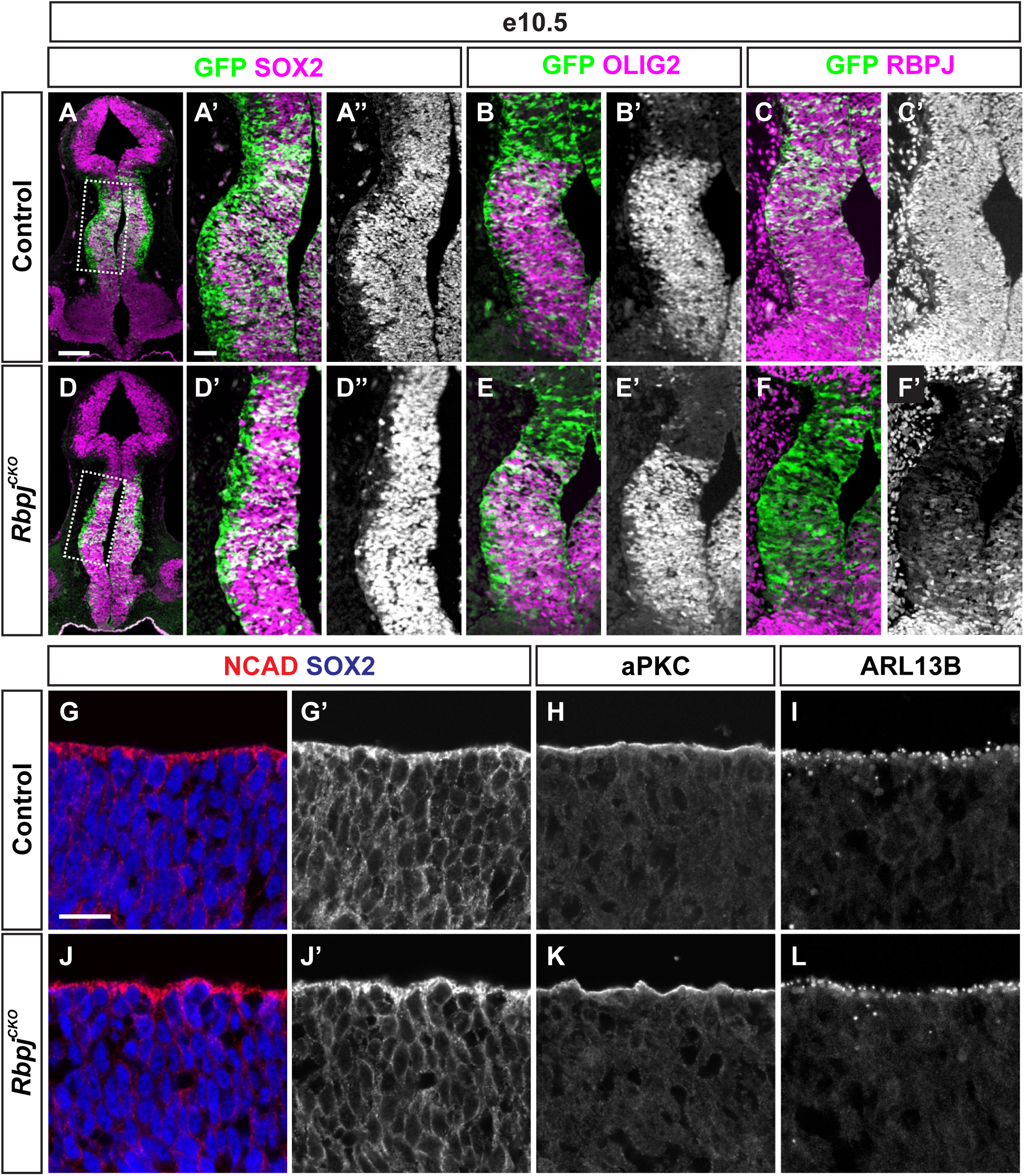
Loss of RBPJ precedes disruption of the VZ. **(A-L)** Coronal sections of e10.5 control and *Rbpj^CKO^* brains. (A-F) Immunofluorescence (IF) for SOX2, OLIG2, RBPJ, and GFP. Dashed white boxes indicate the regions shown at higher magnification. In all the higher-magnification images, the apical surface is oriented to the right. Scale bars: 200 µm (low mag) and 50 µm (high mag). (G-L) High-magnification images of the VZ with IF for NCAD, SOX2, aPKC, and ARL13B. In all images, the apical surface is oriented upward. Scale bars: 20 µm.

**Supplementary Figure S3:**
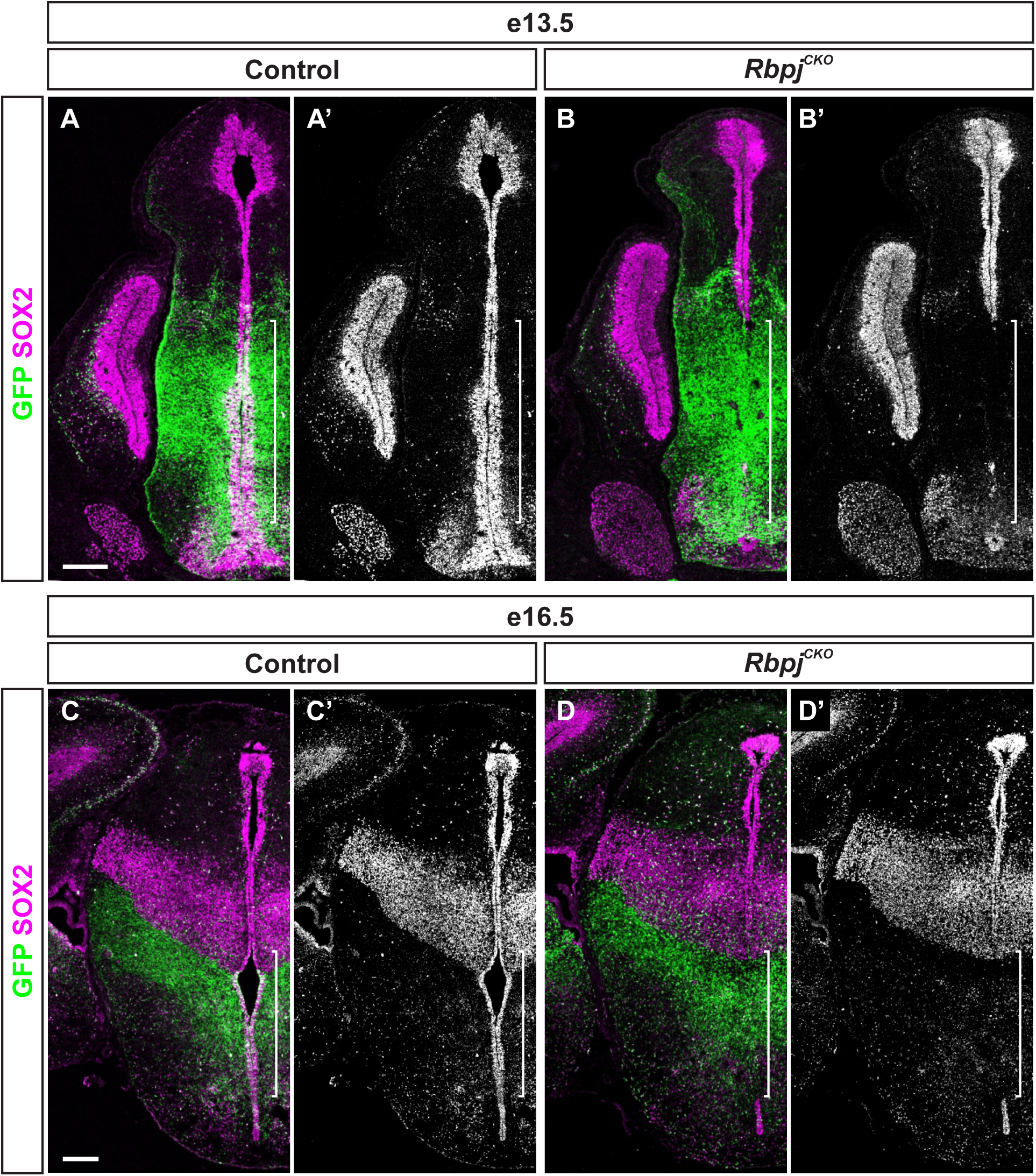
Loss of *Rbpj* results in sustained disruption of the VZ. **(A-D)** Coronal sections through the diencephalon of e13.5 and e16.5 control and *Rbpj^CKO^* brains. Immunofluorescence (IF) images show SOX2 and GFP. Brackets denote the region along the VZ lining the third ventricle that is disrupted in *Rbpj^CKO^* brains. Scale bars: 200 µm.

**Supplementary Figure S4:**
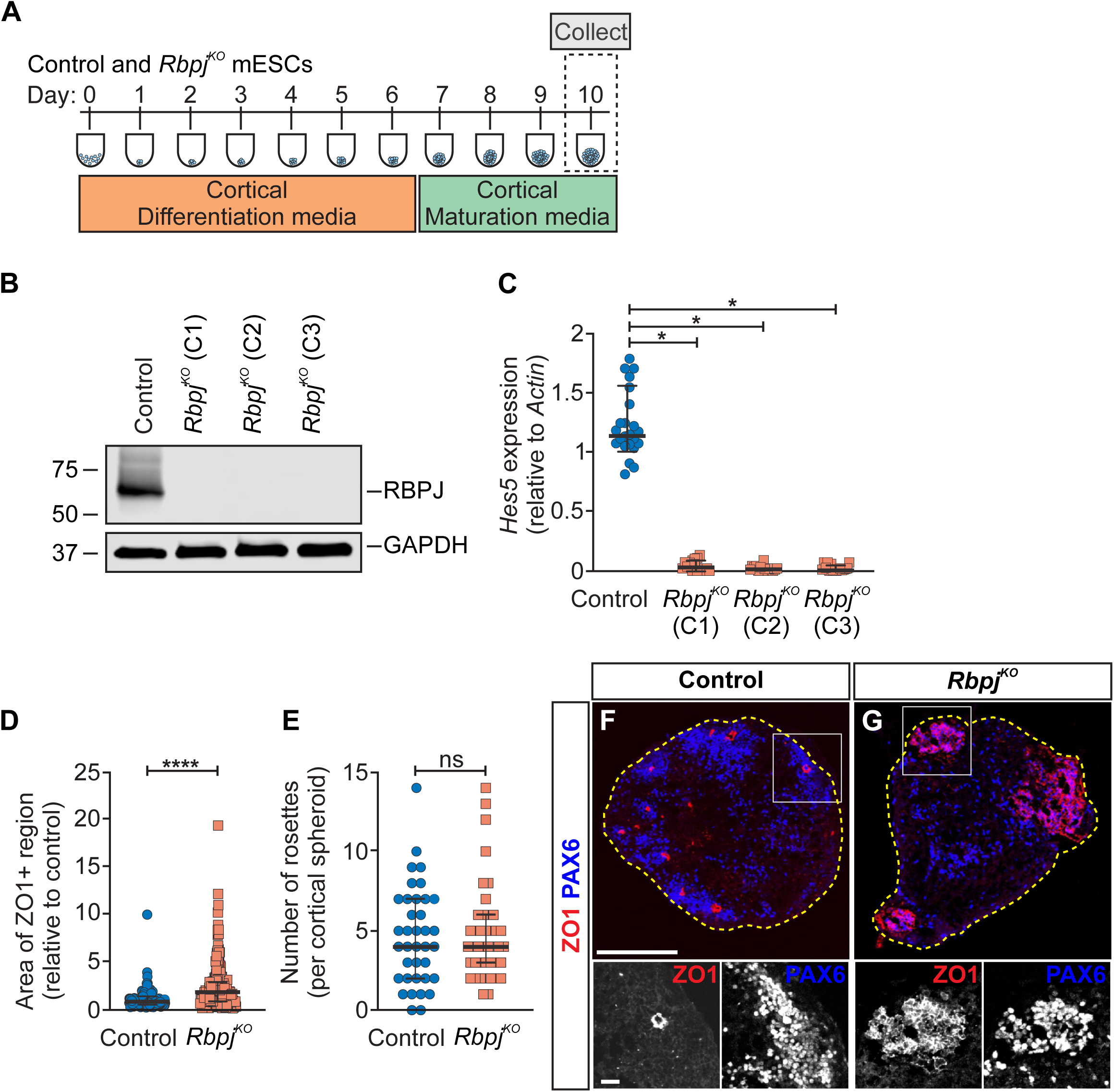
Constitutive loss of Rbpj alters neural rosette organization in cortical spheroids. **(A)** Schematic of the cortical spheroid differentiation protocol. mESCs are transfected with either a non-targeting control or CRISRP-Cas9 guides targeting *Rbpj*. Control or *Rbpj^KO^*mESCs are differentiated into cortical spheroids for 10 days. **(B)** Western blot analysis of RBPJ protein in control and three independently generated *Rbpj^KO^*mESCs clones (C1-C3). GAPDH serves as a loading control. **(C)** *Hes5* expression in control and *Rbpj^KO^* cortical spheroids derived from three independent clones (C1-C3) and normalized to *Actin*. Each data point represents a technical replicate from 4-6 independent experiments, with 3-4 technical replicates per experiment. Bars represent the median with interquartile range. Kruskal-Wallis test, *p = 0.04 (C1), *p=0.04 (C2), and *p=0.01 (C3). **(D)** Quantification of individual ZO1+ region areas. Each data point represents the area of a single ZO1+ region across three independent experiments. Values from each experiment are normalized to the mean ZO1+ area of the corresponding controls to account for experiment-to-experiment variability in spheroid size. Median control: 0.80 (n = 176); median *Rbpj^KO^* = 1.76 (n = 203). Mann-Whitney test, ****p < 0.0001. **(E)** Quantification of ZO1+ neural rosettes. Each data point represents the number of rosettes counted within a single spheroid section collected across three independent experiments. Bars represent the median with interquartile range. Median control: 4 rosettes (n = 37), median *Rbpj^KO^*: 4 rosettes (n = 42). Mann-Whitney test, ****p < 0.0001. **(E-J)** Immunofluorescence (IF) images of control and *Rbpj^KO^*cortical spheroids, stained for ZO1 and PAX6. Yellow dashed lines outline the spheroids. Dashed white boxes indicate the regions shown at higher magnification below. Scale bars: 200 µm (low mag) and 20 µm (high mag).

**Supplementary Figure S5:**
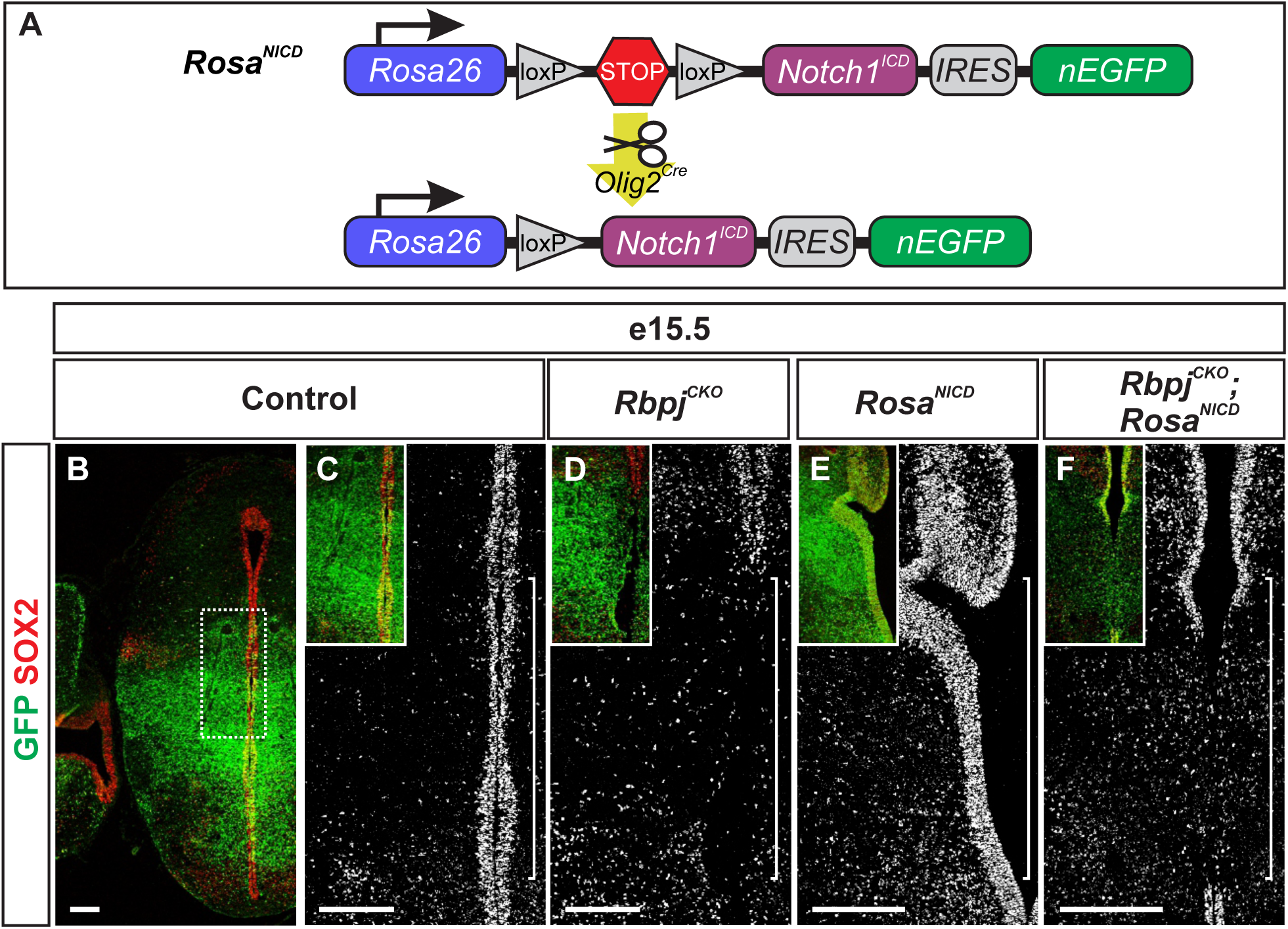
Altered Notch signaling activity disrupts the VZ. **(A)** Schematic of *Olig2^Cre^*-mediated mouse manipulations. In *Rosa^NICD^* mice, *Olig2^Cre^*-mediated recombination removes a floxed stop sequence, resulting in expression of *Notch1* intracellular domain (NICD) and consequently increased Notch signaling activity. **(B-F)** Coronal sections through the diencephalon of e15.5 control, *Rbpj^CKO^*, *Rosa^NICD^*, and *Rbpj^CKO^;Rosa^NICD^*embryos. Immunofluorescence (IF) for SOX2 and GFP. Brackets denote the region along the VZ lining the third ventricle that is disrupted across genotypes. Scale bars: 200 µm.

## MATERIALS AND METHODS

### Animal preparation and tissue analysis

*Olig2^Cre^* mice (*Olig2^tm1(cre)Tmj^*; RRID:MGI:3774124) were generated as previously described^40^. Cre mice were crossed with *Rosa26^eGFP^* transgenic reporter mice (*B6;129-Gt(ROSA)26Sor^tm2Sho^/J*; Jackson Laboratory Strain #004077; RRID:IMSR_JAX:004077)^41^, *Rbpj* transgenic floxed mice (*Rbpj^tm1Hon^*; RRID:MGI:3583755)^42^, and *Rosa^NICD^* transgenic floxed mice (*Gt(ROSA)26Sor^tm1(Notch1)Dam^/J*; Jackson Laboratory Strain #008159; RRID:IMSR_JAX:008159)^65^. All mice were maintained and tissue collected in accordance with guidelines set forth by the UCLA Institutional Animal Care and Use Committee. Mouse brain tissues were collected and fixed in chilled 4% paraformaldehyde (PFA, Ted Pella Inc, Cat #NC1537886) for the following durations of time: e10.5-e11.5 (1 Hour), e12.5 (2 Hours), e13.5 (3 Hours), e14.5 (4 Hours), e15.5-e16.5 (5 Hours), E17.5-P3 (Overnight).After fixation, the tissue was cryoprotected in 30% sucrose, mounted in Tissue-Tek® optimal cutting temperature (OCT) compound (Sakura Finetek USA, Cat #4583), cryosectioned into 12 µm thick sections on a Leica CM3050S cryostat, and processed for immunohistochemistry or in situ hybridization as previously described^66–68^.

#### Cell culture and creation of clonal knockout cell lines

HM1 mouse embryonic stem cells (mESCs) were used in all experiments. The parental HM1 mESCs were derived from a male mouse. To facilitate binding, mESCs were grown on CellBIND dishes (Corning, Cat # 08757214) coated with 0.1% gelatin (Sigma-Aldrich, Cat # 501785182). For maintenance, mESCs were plated on primary mouse fibroblasts treated with Mitomycin C (Thermo Fisher Scientific Cat #AAJ63193MA). mESCs were cultured in mESC media comprised of DMEM high glucose media (Gibco, Cat # SH30081FS), 15% FBS (Gibco, Cat #A5256701), 1x MEM non-essential amino acids (Gibco, Cat #11140050), 1x penicillin/streptomycin (Gibco, Cat # 15140163), 1x Glutamax (Gibco, Cat #35050061), 1x EmbryoMax® nucleosides (Millipore Sigma, Cat #ES-008-D), 55 mM 2-mercaptoethanol (Gibco, Cat #21985023), and 1000 U/ml ESGRO LIF (Millipore Sigma, Cat # ESG1107).

Clonal knockout mESCs were generated using a dual guide sgRNA strategy. Briefly, two sgRNAs targeting *Rbpj* were designed using Benchling and cloned into pSpCas9(BB)-2A-Puro (PX459, Addgene Cat #48139)^69^. Plasmids were electroporated into mESCs using the Lonza nucleofection system (Nucleofector 2b Device using the program A-023 and Lonza Cell Nucleofector Kit Cat #VAPH-1001). mESCs were cultured under feeder-free conditions in 2i media comprised of Dulbecco’s Modified Eagle’s Medium F12 (Gibco, Cat # 11320082) and Neurobasal Medium (Gibco, Cat #21103049) (1:1 ratio) supplemented with N-2 Supplement (Gibco, Cat #17502048), B-27 Supplement (Gibco, Cat #17504044), 1x penicillin/streptomycin (Gibco, Cat # 15140163), 1x Glutamax (Gibco, Cat #35050061), 40 mg/ml Bovine Serum Albumin (Supelco Inc, Cat #A797950ML), 55 mM 2-mercaptoethanol (Gibco, Cat #21985023), 5 mM CHIR99021 (Axon Medchem LLC, Cat #1386), 1 mM PD98059 (Axon Medchem LLC, Cat #1223), and 1000 U/ml ESGRO LIF (Millipore Sigma, Cat # ESG1107). 24 h post nucleofection, cells were selected with 1.5 mg/ml puromycin (Thermo Fisher Scientific, Cat # BP2956100) for 48 h. One week later, individual mESC colonies were manually picked, expanded, and screened by PCR to detect deletion of the genomic DNA segment between the two guides. Sequence of sgRNAs used: Non-targeting control (NTC)^70^: 5’-AAAAAGTCCGCGATTACGTC-3’; *Rbpj* sgRNA #1: 5’-CACGCCAGTTCACAACAGTG-3’; *Rbpj* sgRNA #2: 5’-AGGAGTTCACAGTTAGAGA-3’; *Rbpj* amplicon sequencing primer Fwd: 5’-CTTTCTTTGTGCGTGCCTCC-3’; *Rbpj* amplicon sequencing primer Rev: 5’-TCAAAGCAGAGTCAAGTCTTACCA-3’. To generate a deletion like the *Rbpj* conditional knockout mouse, the *Rbpj* guides targeted exons 6 and 7.

### Reagents and antibodies

Primary antibodies used for immunohistochemistry were as follows: mouse anti-ARL13B (Antibodies Incorporated, Cat #75-287, RRID:AB_2341543), rabbit anti-βCAT (Millipore Sigma, Cat #C2206, RRID:AB_476831), chicken anti-GFP (Aves Labs, Cat #GFP-1020, RRID:AB_2307313), rabbit anti-NCAD (Abcam, Cat #ab12221, RRID:AB_298943), rabbit anti-NCAD (Clone D4R1H, Cell Signaling Technology, Cat #13116S, RRID:AB_2687616), mouse anti-neurofilament (Clone 2H3, Developmental Studies Hybridoma Bank, Cat #2H3, RRID:AB_531793), guinea pig anti-OLIG2 (RRID: AB_2715520)^71^, rabbit anti-OLIG2 (Millipore Sigma, Cat #AB9610, RRID:AB_570666), rabbit anti-PKCζ (Clone C-20, Santa Cruz Biotechnology, Cat #sc-216, RRID:AB_2300359), rat anti-RBPJ (Clone 1F1, Active Motif, Cat #61506, RRID:AB_2614972), rabbit anti-RBPSUH (Clone D10A4, Cell Signaling Technology, Cat #5313T, RRID:AB_266555), goat anti-SOX2 (Clone Y-17, Santa Cruz Biotechnology, Cat #sc-17320, RRID:AB_2286684), mouse anti-TUBA1 (Clone 6-11B-1, Sigma Aldrich, Cat #T6793, RRID:AB_477585), mouse anti-TUBB3 (Clone TUJ1, previously Covance Cat #MMS-435P and currently BioLegend Cat #801202, RRID:AB_2313773). All antibodies were diluted to a working concentration in Antibody Block solution comprised of 1% heat-inactivated horse serum (Cytiva Hyclone™, Cat # SH3007403) in PBS (Alkali Scientific, Cat #RB1200) with 0.1% Triton X-100 (Thermo Fisher Scientific, Cat #BP151-500). Secondary antibodies conjugated to IRDye® infrared dyes for Western blot or Alexa Fluor dyes for immunofluorescence were purchased from LI-COR Biosciences, Thermo Fisher Scientific, and Jackson ImmunoResearch Laboratories. RNA probes were generated by in vitro transcription of PCR products amplified from mouse spinal cord cDNA. Primers against the following genes were designed using the program Primer3. *Hes5*, forward 5’-GGATGAGCTCGTTCCTCTGG-3’ and reverse 5’-GAG<u>ATTAACCCTCACTAAAGGGA</u>GCAGGCTGAGTGCTTTCCTA-3’. The underlined text denotes a T3 polymerase binding site incorporated into the reverse primer.

### Evans blue dye injections

P0-5.5 neonatal mice were euthanized in accordance with guidelines set forth by the UCLA Institutional Animal Care and Use Committee. The skin was removed from the skull and, with the skull still intact, a glass capillary needle was used to slowly inject 5-10 µl of 4% Evans blue dye (Thermo Fisher Scientific, Cat #A16774.18) diluted in sterile PBS into the lateral ventricles to visualize the pathway of the cerebrospinal fluid (CSF) through the ventricular system^72,73^. After Evans blue dye addition, the brain was removed from the skull, briefly rinsed in PBS, and then fixed overnight in 4% PFA. After fixation, the brains were rinsed thoroughly in PBS and then assessed for obstructive hydrocephalus. Brains were then scored as either “normal” or “obstructed” based on the distance the dye traveled.

### Vibratome sections

The brains from the Evans blue dye assay were sectioned to better assess where the obstruction occurred in the P0-5.5 neonatal mutant mice. After Evans blue dye injection into the lateral ventricles, the brains were fixed, thoroughly rinsed in PBS, mounted in 4% low melting agarose prepared in PBS, and then 300 µm serial coronal sections were cut on a Leica VT1000S vibratome.

### Tissue clearing

To better see where the obstruction occurred in the P5.5 neonatal mutant mouse brains, after Evans blue dye was injected into the lateral ventricles the brains were made transparent as previously described^74^. The protocol was modified slightly from what was reported. Briefly, the tissue was fixed in 4% PFA overnight at 4°C, rinsed thoroughly in PBS, bleached overnight in a solution of 33.3% H2O2/13.3% DMSO/53.4% MeOH at 4°C, dehydrated in a series of MeOH washes, post-fixed overnight in a solution of 20% DMSO/80% MeOH at 4°C, rinsed in MeOH, and then cleared in BABB comprise of 33.3% benzyl alcohol (Thermo Fisher Scientific, Cat #AA41218AK) and 66.6% benzyl benzoate (Millipore Sigma, Cat #501854457).

#### Western Blotting

Whole cell extracts from NIH/3T3 cells and NPCs were prepared in RIPA lysis buffer comprised of 50 mM Tris-HCl pH-7.4 (Gold Biotechnology, Cat #T095100), 150 mM NaCl (Thermo Fisher Scientific, Cat #S271), 2% NP-40 (United States Biological, Cat #N3500), 0.25% Sodium Deoxycholate (Sigma Aldrich Fine Chemicals Biosciences, Cat #D6750100G), 0.1% SDS (Thermo Fisher Scientific, Cat #AAJ1822036), 1mM DTT (Gold Biotechnology, Cat #DTT100), 10% glycerol (Thermo Fisher Scientific, Cat #G331), and 1x SIGMAFAST™ protease inhibitor cocktail (Millipore Sigma, Cat #S8830). Samples were resuspended in 4x NuPAGE-LDS sample buffer (Thermo Fisher Scientific, Cat #NP0007) or 4x Protein Sample Loading Buffer (LI-COR Biosciences, Cat #928-40004), incubated at 37°C for 30 min, and subjected to SDS-PAGE. The resolved proteins were transferred onto a PVDF membrane (Millipore Sigma, Cat #IPVH00010) using a wet electroblotting system (Nupage system, Thermo Fisher Scientific) followed by immunoblotting.

### Cortical spheroid protocol

Mouse embryonic stem cells (mESCs) were plated on Nuclon™ Sphera™ 96-well U-shaped-bottom microplates (Thermo Fisher Scientific, Cat #12566430) with 300,000 cells/well in cortical differentiation media. The cortical differentiation media consisted of DMEM high glucose media (Gibco, Cat # SH30081FS) supplemented KnockOut™ Serum Replacement (Gibco, Cat # 10828028), 1mM sodium pyruvate (Gibco, Cat #11360070), 1x MEM non-essential amino acids (Gibco, Cat #11140050), and 1x 2-mercaptoethanol (Gibco, Cat #21985023). ROCK inhibitor (Y-27632, BD Biosciences, Cat #BDB562822) was added at 10uL/mL of cortical differentiation media only on day of plating (Day 0). 100uL of media described above was added to each well the next day (Day 1). For days 2-5, half of the media was aspirated off and replaced with 100uL of fresh differentiation media. On day 6, Notch inhibitor 10 nM StemSelect Compound E (Millipore Sigma, Cat #5305090001) was added to the prepared differentiation media. On day 7, all media was aspirated out of each well and replaced with 150uL of cortical maturation media. The cortical maturation media consisted of Dulbecco’s Modified Eagle’s Medium F12 (Gibco, Cat # 11320082) supplemented with N-2 Supplement (Gibco, Cat #17502048), 1x GlutaMAX (Gibco, Cat #35050061), and 1x penicillin/streptomycin (Gibco, Cat # 15140163). For days 8 and 9, half media change was done as described in days 2-5 with cortical maturation media. On day 10, the spheroids were processed for immunofluorescence or qPCR. For immunofluorescence, the spheroids were fixed for 30 minutes in 4% PFA followed by thorough rinsing in PBS before being mounted in a 30% sucrose solution.

### Image analysis

Fluorescence images were collected using an LSM5 Exciter confocal imaging system, Nikon eclipse Ti2 widefield microscope, and Nikon A1R HD25 laser scanning confocal microscope. Images were collected and processed using LSM exciter, Zeiss Efficient Navigation (ZEN), Nikon Imaging Software (NIS) Elements, Fiji^75^, and Adobe Photoshop software.

### Statistical Analysis

Graphing and statistical analyses were performed using Prism version 10 (GraphPad Software). Significance was assumed when *p < 0.05, **p < 0.01, ***p < 0.001. ns, p > 0.05.

### Additional software and AI tools use

The manuscript was written with grammatical assistance from Grammarly (Superhuman Platform Inc). Citations were generated using Paperpile (Paperpile LLC). After using these services, the authors reviewed and edited the content as needed and take full responsibility for the content of the publication.

